# Synaptic vesicle glycoprotein 2C (SV2C) modulates dopamine release and is disrupted in Parkinson’s disease

**DOI:** 10.1101/077586

**Authors:** Amy R. Dunn, Kristen A. Stout, Minagi Ozawa, kelly M. Lohr, Alison I. Bernstein, Yingjie Li, Minzheng Wang, Carmelo Sgobio, Namratha Sastry, Huaibin Cai, W. Michael Caudle, Gary W. Miller

**Affiliations:** Rollins School of Public Health, Emory University, Atlanta, GA 30322.; Center for Neurodegenerative Disease, Emory University, Atlanta, GA 30322.; Transgenics Section, Laboratory of Neurogenetics, National Institute on Aging, National Institutes of Health, Bethesda, MD 20892.

**Keywords:** Parkinson’s disease, synaptic vesicles, dopamine, SV2C

## Abstract

The synaptic vesicle glycoprotein 2 (SV2) family of proteins are involved in synaptic function throughout the brain. The ubiquitously expressed SV2A has been widely implicated in epilepsy, though SV2C with its restricted basal ganglia distribution has no known function. SV2C is emerging as a potentially relevant protein in Parkinson’s disease, as it is a genetic modifier of nicotine neuroprotection and sensitivity to L-DOPA. Here we identify SV2C as a mediator of dopamine homeostasis and report that disrupted expression of SV2C within the basal ganglia is a pathological feature of Parkinson’s disease (PD). Genetic deletion of SV2C leads to reduced dopamine release in the dorsal striatum as measured by fast-scan cyclic voltammetry, reduced striatal dopamine content, disrupted alpha-synuclein expression, deficits in motor function, and alterations in neurochemical effects of nicotine. Further, SV2C expression is dramatically altered in postmortem brain tissue from PD cases, but not in Alzheimer’s disease, progressive supranuclear palsy or multiple system atrophy. This disruption was paralleled in mice overexpressing mutated α-synuclein. These data establish SV2C as a novel mediator of dopamine neuron function and suggest that SV2C disruption is a unique feature of PD that likely contributes to dopaminergic dysfunction

## INTRODUCTION

Synaptic vesicles, particularly those within the dopaminergic nigrostriatal pathway, have two important roles: packaging transmitter for neurotransmission and sequestering compounds that may have adverse effects on the cell, such as cytosolic dopamine(1–3). Ineffective sequestration of dopamine leads to progressive cell loss and heightened vulnerability to dopaminergic toxicants in mice(4–10). Mutations in the gene encoding the vesicular monoamine transporter 2 (VMAT2) lead to infantile parkinsonism(11), while increased expression of VMAT2 is associated with decreased risk for Parkinson’s disease (PD)(12, 13). Dopamine vesicle function is impaired in patients with PD, suggesting that dysfunctional dopamine handling is fundamental to the disease(14). Genetic mutations in several vesicular-associated proteins, such as α-synuclein, LRRK2 and synaptojanin, can lead to disrupted presynaptic dopamine handling and are commonly linked to PD (15–22).Disrupted vesicle function may represent a common pathway to degeneration and identifying novel mediators of vesicular function could provide insight to our understanding of disease pathogenesis.

Proteins within the synaptic vesicle glycoprotein 2 (SV2) family are thought to positively modulate vesicular function in a variety of ways, possibly by aiding in vesicular trafficking and exocytosis, interacting with synaptotagmin-1 or facilitating vesicular transmitter uptake(23–33). SV2A has been strongly implicated in epilepsy and is the specific pharmacological target for the antiepileptic levetiracetam; however, despite extensive research the molecular function of SV2 proteins has not been fully described. SV2C is distinguished from SV2A and SV2B by its enriched expression in the basal ganglia and preferential localization to dopaminergic cells in mice(34, 35). Intriguingly, SV2C was recently identified as a genetic mediator of one of the most robust environmental modulators of PD risk: nicotine use, which is strongly protective against PD (36, 37). Variation within the SV2C gene was also found to predict PD patient sensitivity to L-DOPA (43). These data suggest that SV2C may play a particularly important role in the basal ganglia, though there has been no experimental evidence linking SV2C to dopaminergic function, neurochemical effects of nicotine, or Parkinson’s disease.

#### Significance

Here we describe a role for the synaptic vesicle glycoprotein 2C (SV2C) in dopamine neurotransmission and Parkinson’s disease (PD). SV2C is expressed on the vesicles of dopamine-producing neurons and genetic deletion of SV2C causes a reduction in synaptic release of dopamine. The reduced dopamine release is associated with a decrease in motor activity. SV2C is suspected of mediating the neuroprotective effects of nicotine and we show an ablated neurochemical response to nicotine in SV2C-knockout mice. Lastly, we demonstrate that SV2C expression is specifically disrupted in mice that express mutated alpha-synuclein and in humans with PD. Together, these data establish SV2C as an important mediator of dopamine homeostasis and a potential contributor to PD pathogenesis.

We first characterized SV2C expression in multiple mouse models of PD in order to establish potential alterations in SV2C following dopaminergic cell loss. To directly test the hypothesis that SV2C is involved in dopamine function within the basal ganglia, we then developed a line of mice lacking SV2C (SV2C-KO). We quantified striatal dopamine content and dopamine metabolites in SV2C-KO animals with high performance liquid chromatography (HPLC). We then evaluated alterations in dopamine- and PD-related motor behavior and protein expression following genetic deletion of SV2C and and evaluated a possible interaction between SV2C and α-synuclein. Next, we performed fast-scan cyclic voltammetry to measure stimulated dopamine release in the dorsal striatum of SV2C-KO and wildtype (WT) mice at baseline and in the presence of nicotine. Finally, we analyzed SV2C expression in the basal ganglia of postmortem PD cases and other neurodegenerative diseases. The data described below establish SV2C as a mediator of dopamine dynamics and provide a functional basis for a role for SV2C in PD.

## RESULTS

### Distribution of SV2C in mouse basal ganglia

There is no acceptable SV2C antibody currently commercially available; as such, we designed two polyclonal antibodies raised against amino acids 97–114 of (1) mouse SV2C (mSV2C; antibody = mSV2CpAb) and (2) human SV2C (hSV2C; antibody = hSV2CpAb). To confirm the specificity of the hSV2CpAb, we performed immunoblots on transfected HEK293 cells overexpressing either hSV2A or hSV2C. The hSV2CpAb immunoreactivity was present in hSV2C-transfected cell lysate and not in hSV2A-transfected cell lysate. Preincubating the hSV2CpAb with the immunizing peptide (“antigen-blocking”) ablates immunoreactivity (Fig. 1A). We performed shRNA knockdown of SV2C in Neuro-2a (N2a) cells, which are mouse-derived and endogenously express SV2C but not SV2A or SV2B(38). Two sequences of shRNA targeting SV2C mRNA were designed, along with a scrambled shRNA sequence (OriGene). SV2C shRNA robustly reduced normal SV2C immunoreactivity in N2a cell lysates (Fig. 1C).

**Figure 1.**
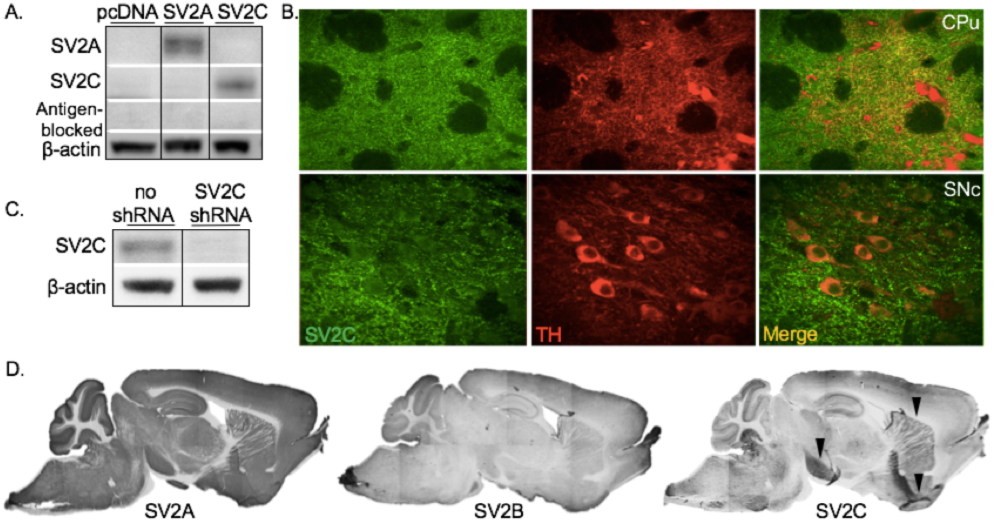
Polyclonal SV2C antibody characterization. We designed polyclonal SV2C antibodies. (A) hSV2C antibody recognizes SV2C but does not recognize SV2A in transfected HEK293 cells. Pre-incubating the hSV2C antibody with the immunizing peptide (“antigen-blocking”) eliminates antibody reactivity. (B) SV2C is highly colocalized with dopaminergic marker tyrosine hydroxylase (TH) in the substantia nigra pars compacta (SNc) and dorsal striatum (CPu) of mouse. (C) Knocking down endogenous SV2C in Neuro-2a cells (which have no endogenous SV2A or SV2B) using RNAi reduces mSV2C antibody reactivity. (D) Sagittal 3–3’ diaminobenzidine-(DAB) stained sections showing ubiquitous SV2A and SV2B expression compared to SV2C expression enriched in the basal ganglia using the mSV2C antibody. Black arrows indicate areas of highest SV2C staining in the mouse brain: midbrain, dorsal striatum and ventral pallidum. *Immunoblots separated by black lines indicate bands that came from the same blot and were cropped for easy comparison between lanes*.

Staining patterns observed with our mSV2CpAb were consistent with published antibodies(34, 35). SV2A and SV2B are strongly expressed throughout the mouse brain, whereas SV2C is preferentially expressed in limited nuclei (Fig. 1D). We identified strong SV2C immunoreactivity in the ventral pallidum (VP), as well as in cell bodies in the midbrain (substantia nigra pars compacta, SNpc; ventral tegmental area, VTA). SV2C-positive fibers were additionally observed in the substantia nigra pars reticulata (SNpr), dorsal striatum (CPu) and nucleus accumbens (NAc). SV2C was highly colocalized with dopaminergic marker tyrosine hydroxylase (TH) in the CPu and substantia nigra (Fig. 1B).

### SV2C expression in animal models of dopamine degeneration

Next, we explored SV2C expression in mouse models of dopamine cell loss. Each model recapitulates a distinct pathogenic process of cell loss associated with PD, allowing us to evaluate potential involvement of SV2C in various mechanisms of PD-related degeneration.

#### 1-methyl-4-phenyl-1,2,3,6-tetrahydropyridine(MPTP) intoxication in mice

In order to investigate whether SV2C expression is altered following a lesion to the nigrostriatal tract, we administered three different dosing paradigms of MPTP to animals to model various stages of degeneration. A 2x15 mg/kg dose, which typically induces a mild loss of striatal dopamine terminals, resulted in a 52% loss of striatal TH immunoreactivity, as measured by immunoblotting (Saline: 10700996AU, MPTP: 5010330AU, p<0.01). A dose of 4x15mg/kg, which results in an intermediate loss of both dopamine terminals and SNpc cell bodies, induced a 68% loss of striatal TH. A 5x20 mg/kg intoxication, a dose that typically results in significant SNpc cell loss, ablated 86% of striatal TH (Saline: 6310104AU, MPTP: 91691.3AU, p<0.0001). Concordant with dopamine terminal loss, striatal SV2C levels were reduced after MPTP (n=6, p<0.05 for the 4x15mg/kg paradigm, Fig 2A). SV2C expression patterns appear unaltered after MPTP at any dose

#### VMAT2-LO model of Parkinson’s disease

To determine whether long-term dysregulation of dopamine storage results in altered expression of SV2C, we evaluated SV2C in animals with a 95% reduction in VMAT2 expression (VMAT2-LO). This genetic reduction of VMAT2 results in reduced vesicular storage of dopamine, progressive loss of dopaminergic cells and resultant motor and nonmotor impairments(5, 10, 39). Similar impairment of VMAT2 function is seen in idiopathic PD(14). As expected, in aged (20–24 month) VMAT2-LO mice, we observed a 53% loss of striatal DAT (Fig. 2B). This terminal loss is consistant with an observed slight but statistically insignificant reduction in striatal SV2C, with no change in expression pattern

**Figure 2.**
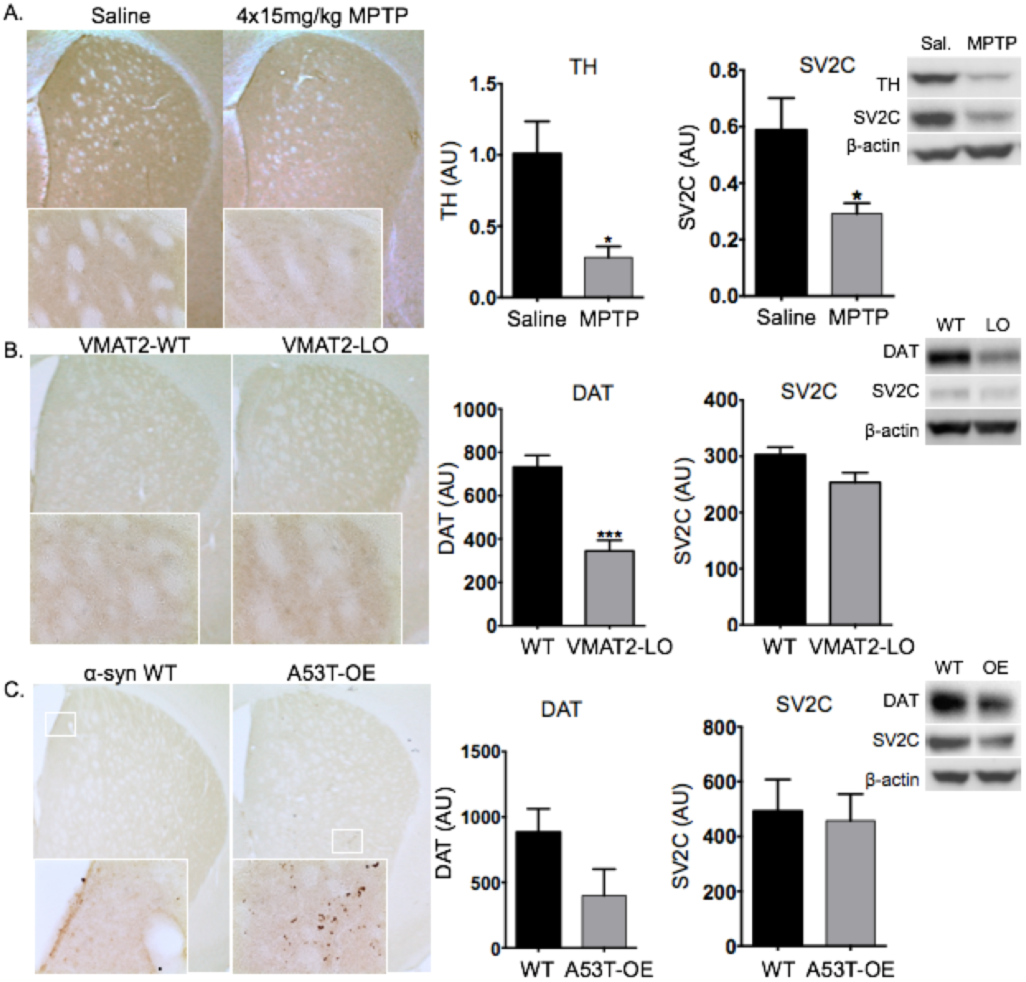
SV2C expression in mouse models of PD. (A) Striatal TH is 68% reduced after a 4x15mg/kg dose of MPTP (n=6, p<0.01). Striatal SV2C is reduced after MPTP (p<0.05). Representative immunohistochemical staining after MPTP reveals that striatal SV2C expression patterns are unchanged.(B) Striatal SV2C levels are slightly, but not significantly, reduced in aged VMAT2-LO animals. SV2C expression patterns remain unaltered, as shown by representative immunohistochemistry in the dorsal striatum. (C) An increase of SV2C-positive punctate staining is observed in the striatum of mouse overexpressing A53T α-synuclein (A53T-OE) as compared to WT mice, particularly in the periventricular mediodorsal striatum. Dense clusters of punctate SV2C are found distributed sparsely in the striatum of A53T-OE mice.

#### A53T α-synuclein overexpression in mice

To determine if α-synuclein disruption alters SV2C, we explored SV2C expression in mice overexpressing PD-associated A53T α-synuclein (A53T-OE) under control of the *Pitx3* promoter, limiting overexpression to midbrain dopamine neurons. These mice display impaired vesicular dopamine release, progressive dopamine terminal degeneration and subsequent motor impairment by 12 months of age(40). Overexpression of A53T α-synuclein in aged animals (12mo; 1 male, 3 female) resulted in an increase in punctate SV2C staining observed in the striatum as compared to WT littermates (Fig. 2C), indicating that SV2C expression patterns are altered with overexpression of mutated α-synuclein. This punctate staining was particularly strong in the periventricular region of the dorsomedial striatum and sparse with dense clusters of puncta throughout the dorsolateral striatum (Fig 2C, right panel)

### Genetic deletion of SV2C reduces striatal dopamine without overt changes in related proteins or dopamine metabolism

To determine a potential role of SV2C in dopamine-related neurochemistry and behavior, we generated a line of SV2C-KO mice (Fig. 3A–B). We performed HPLC in dorsal striatal tissue of SV2C-KO and WT animals to determine if genetic deletion of SV2C altered dopamine tone or metabolism. SV2C-KO animals had a 33% reduction in striatal dopamine content (WT: 741.8±53.6ng/mL, KO: 496.8±53.8ng/mL, p<0.01, n=7, Fig 3D), a 19% reduction in DOPAC (WT: 113.5±5.5ng/mL, KO: 91.85±6.5ng/mL, n=7, p<0.05, Fig 3E), a dopamine metabolite, and no difference in DOPAC:dopamine ratio (not shown). This reduction in dopamine content does not appear to be the result of reduced tyrosine hydroxylase (TH), the rate-limiting enzyme in dopamine synthesis, and does not result in up- or down-regulated dopamine transporter (DAT) by immunoblot. Further, SV2C deletion does not appear to induce a compensatory upregulation in levels of either SV2A or SV2B (Fig. 2D).

**Figure 3.**
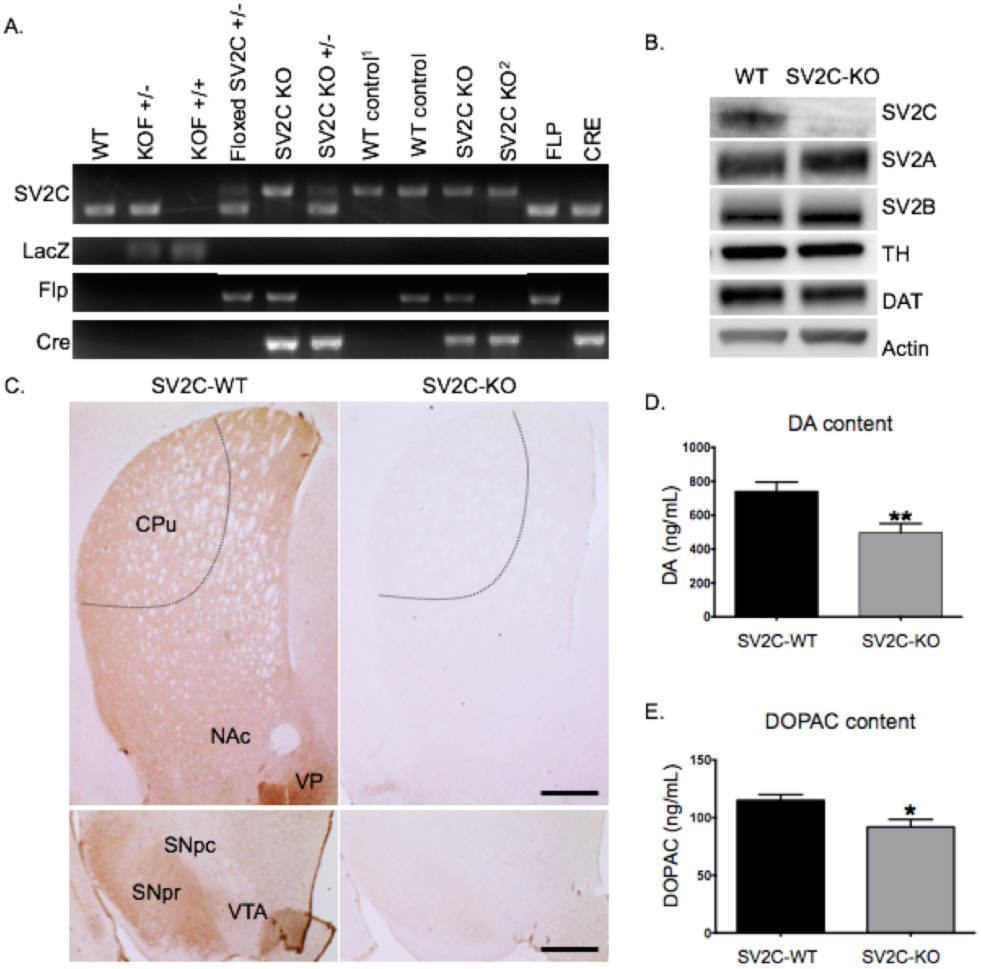
Neurochemical characterization of SV2C-KO mice. (A) PCR genotyping of SV2C-KO mice generated using the EUCOMM “knockout-first” allele. This construct allowed us to generate several useful lines of mice: a preliminary knockout animal with a cassette creating a frt-flanked gene trap inserted into the second exon (KOF), a line with a floxed exon after breeding KOF animals with a flp-recombinase+ line, and finally SV2C-KO mice following a cross with a nestin-driven Cre-recombinase+ line. Animals used for experiments are denoted by a superscript 1 (WT control) and 2 (SV2C-KO). See methods for primer sequences. (B) SV2C-KO does not result in altered expression of either SV2A or SV2B, nor of dopaminergic markers tyrosine hydroxylase (TH) or dopamine transporter (DAT). (C) Genetic deletion of SV2C ablates mSV2C antibody reactivity in the striatum (CPu) and midbrain. Dotted line delineates dorsolateral striatum where electrochemical recordings were taken. (D-E) SV2C-KO results in a 33% reduction of dopamine content in the dorsal striatum (n=7, p<0. 01) and a 19% reduction in the dopamine metabolite DOPAC.

### An interaction between SV2C and α-synuclein

To explore differences in PD-related protein expression in SV2C-KO animals, we performed immunoblotting and immunohistochemistry. SV2C-KO animals had slightly reduced monomeric (~15kD) α-synuclein (69.1±6.17% of WT, n=4–6, p=0.06) and significantly increased high molecular weight (multimeric, ~90kD) α-synuclein expression (3020±561% of WT, n=4–6, p<0.01, Fig. 4B–D). Following the observation of disrupted α-synuclein in SV2C-KO mice along with disrupted SV2C in A53T-OE mice, we performed immunoprecipitations and immunoblotting to investigate an interaction between SV2C and α-synuclein. In striatal preparations from tissue of WT C57BL/6 mice, α-synuclein co-immunoprecipitated with SV2C. As expected, neither TH nor DAT co-immunoprecipitated with SV2C (negative control). Synaptotagmin-1, which has an SV2C binding site(26, 27), did co-immunoprecipitate (positive control, Fig. 4A).

**Figure 4.**
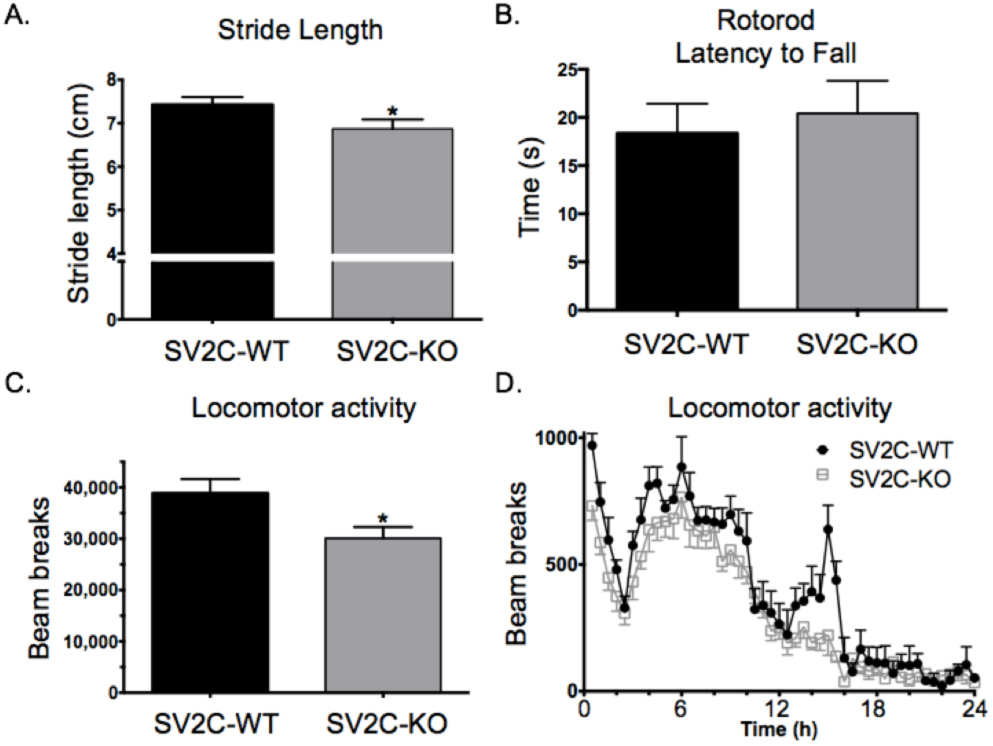
An association between SV2C and α-synuclein. (A) α-synuclein co-immunoprecipitates with SV2C in WT mouse striatal synapotosomal preparations. TH, a cytosolic protein, and DAT, a membrane-associated protein, were included as negative controls. Synaptotagmin-1, which binds to SV2C, was used as a positive control. (B-D) Midbrain homogenates from SV2C-KO animals have a 31% decrease in monomeric (15kD) α-synuclein (n=4–6, p=0.06) and a 30-fold increase in multimeric (90kD) α-synuclein (n=4–6, p<0.01).

### SV2C-KO animals have mild motor deficits

To determine if SV2C-KO results in alterations in motor behavior consistent with their decreased dopamine content, we performed multiple motor function assays. We performed a 24-hour locomotor activity assay, gait analysis and a rotorod test and found that SV2C-KO was associated with mild motor deficits. SV2C-KO animals exhibited reduced locomotor activity, with a 29% reduction in total ambulations over 24 hours (WT: 38900±2770 ambulations, KO: 30100±2200 ambulations, n=7–8, p<0.05). Additionally, SV2C-KO mice displayed a shorter stride length (WT: 7.42±0.17cm, KO: 6.87±0.22cm, n=22–24, p<0.05). SV2C-KO animals showed no differences on latency to fall during the rotorod test (WT: 18.3±3.01sec, KO: 20.4±3.36sec, n=7–8, p=0.52) (Fig. 5). Additionally, SV2C-KO animals had a lower average body size (WT: 19.06±0.91g, KO: 17.24±0.99g, n=53, p<0.05).

**Figure 5.**
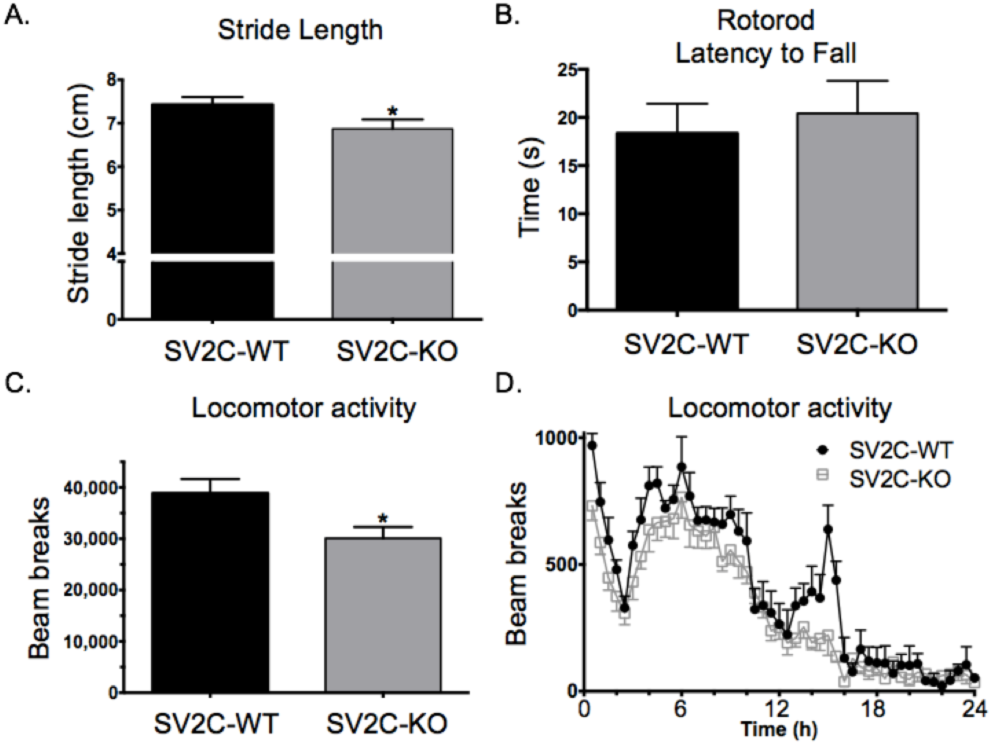
Altered motor behavior of SV2C-KO mice. (A) SV2C-KO animals have about a 10% reduction in stride length as measured by a gait analysis assay (p<0.05). (B) Genetic deletion of SV2C does not result in impairment on the rotorod test. (C) SV2C-KO animals display a 23% reduction in total ambulations in a 24-hour locomotor activity monitoring (n=7–8, p<0.05). (D) This reduction in locomotor activity is apparently driven by an ablation of a circadian peak in activity at the end of the active period.

### Genetic deletion of SV2C results in reduced striatal dopamine release

In order to determine if the observed behavioral deficits of SV2C-KO mice resulted from a reduction in striatal dopamine release, we performed *ex vivo* fast-scan cyclic voltammetry in the dorsolateral striatum of SV2C-KO animals (see Fig. 3B for delineation of dorsolateral striatum). Genetic deletion of SV2C led to a 32% decrease in dopamine release as compared to WT animals (WT: 5.47±0.35µM, KO: 3.71±0.57µM, p<0.05, n=9, Fig. 6).DAT-mediated dopamine clearance rate was enhanced in SV2C-KO animals as evidenced by reduced tau (WT: 0.39±0.03sec, KO: 0.29±0.02sec, p<0.05).

**Figure 6.**
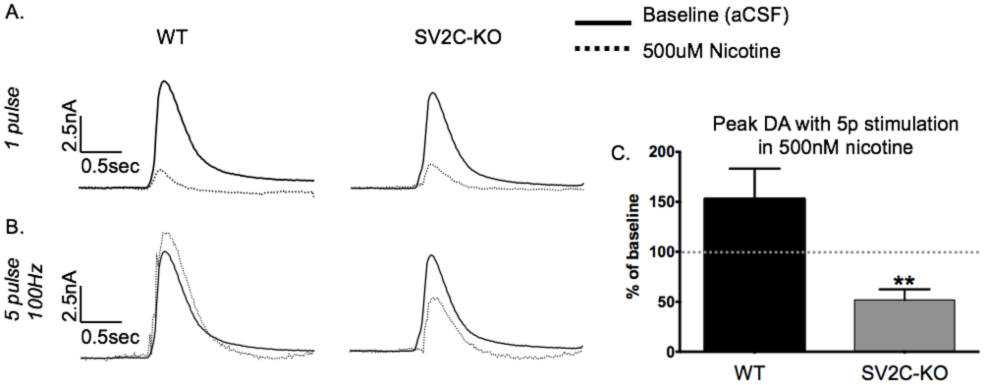
Electrochemical measurement of stimulated dopamine release in SV2C-KO. Genetically ablating SV2C reduces electrically evoked dopamine release by 32% as shown by (A) a representative color blot and (B) quantified over all recording sites in each animal (n=9, p=0.01) and (C) represented by respective current traces.

### SV2C mediates a neurochemical effect of nicotine

In order to assess a potential functional interaction between SV2C and nicotine, we performed FSCV in the presence of 500nM nicotine. This concentration of nicotine typically dampens dopamine release elicited by a single pulse stimulation while enhancing dopamine release elicited by a high intensity (e.g., five pulses at 100Hz) stimulation(41, 42). Genetic deletion of SV2C ablated this effect. In WT tissue, 500nM nicotine enhanced dopamine release elicited by five pulses at 100Hz to 153% of baseline; in SV2C-KO tissue 500nM nicotine *reduced* dopamine release elicited by five pulses at 100Hz to 48% of baseline (p<0.01). These data are presented as current traces (7A–B) and peak dopamine release as a percentage of baseline (single pulse stimulation) (7C).

**Figure 7.**
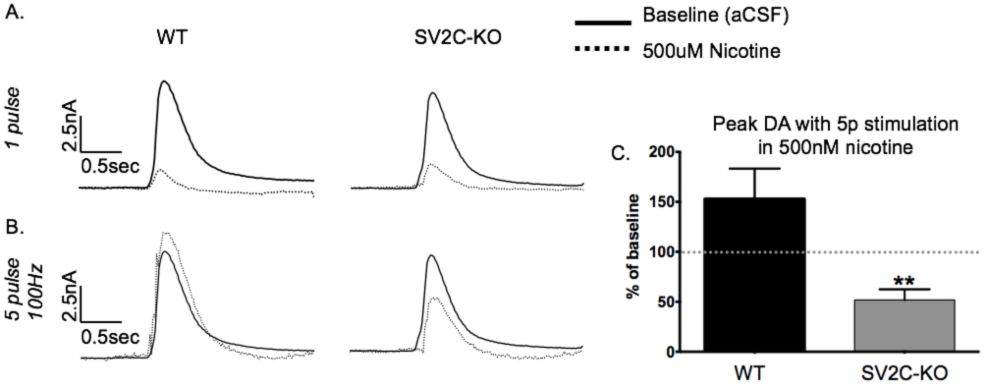
Altered neurochemical effect of nicotine after genetic ablation of SV2C. (A) *Top panels*: Current traces showing dopamine release at baseline and in the presence of 500nM nicotine. Nicotine normally reduces dopamine release elicited by a one-pulse stimulation to about 10% of baseline in both WT and SV2C-KO striatum. *Bottom panels*: Five-pulse stimulations in the presence of nicotine normally increase dopamine release over baseline, but this effect is not seen in SV2C-KO animals. (C) Dopamine release elicited by a five-pulse stimulation in the presence of nicotine represented as percent of baseline (p<0.01).

### SV2C expression in human basal ganglia in control and disease cases

To explore a potential involvement of SV2C in human PD, we obtained human striatum and midbrain samples from brain banks at Emory University and the University of Washington. Control and disease cases were matched for age (72.4±3.9 vs. 71.7±2.3 years, respectively) and sex (71% vs. 68% male, respectively). Four PD cases, three comorbid dementia with Lewy Bodies (DLB)/PD cases, seven Alzheimer’s disease (AD) cases, three multiple system atrophy (MSA, including one with comorbid DLB and one with comorbid olivopontocerebellar atrophy (OPC)), two progressive supranuclear palsy (PSP) cases, and seven age-matched controls were examined (Table 1). MSA and PSP were chosen for their clinical and pathological similarities to PD, and AD was chosen as a non-basal ganglia neurodegenerative disease.

**Table 1.** Descriptions of human control and disease cases.

| Case | Neuropath. Diagnosis | Sex | Age (years) | Illness duration (years) | SV2C pathology |
| --- | --- | --- | --- | --- | --- |
| **Controls** |  |  |  |  |  |
| 1 | -- | M | 72 | -- | None |
| 2 | -- | F | 80 | -- | None |
| 3 | -- | M | 78 | -- | None |
| 4 | -- | F | 88 | -- | None |
| 5 | -- | M | 61 | -- | None |
| 6 | -- | M | 69 | -- | None |
| 7 | -- | M | 59 | -- | ++ |
| **Parkinson's disease** |  |  |  |  |  |
| 8 | PD | M | 78 | 32 | STR: +++ |
| 9 | PD | M | 64 | 12 | STR: +++ |
| 10 | PD | M | 81 | 10 | STR: ++/<br>SN: +++ |
| 11 | PD | M | 67 | 7 | STR: ++/<br>SN: ++ |
| 12 | DLB/PD | M | 68 | 21 | STR: ++ |
| 13 | DLB/PD | F | 62 | 4 | STR: + |
| 14 | DLB/PD | M | 72 | 11 | STR: +/<br>SN: + |
| **Alzheimer's disease** |  |  |  |  |  |
| 15 | AD | M | 63 | Unk | None |
| 16 | AD | M | 75 | Unk | STR: ++ |
| 17 | AD | F | 81 | Unk | None |
| 18 | AD | M | 50 | Unk | None |
| 19 | AD | F | 87 | Unk | None |
| 20 | AD | F | 59 | Unk | None |
| 21 | AD | M | 79 | Unk | None |
| **Multiple system atrophy / Progressive supranuclear palsy** |  |  |  |  |  |
| 22 | MSA | M | 61 | Unk | None |
| 23 | MSA/OPC | F | 80 | Unk | None |
| 24 | MSA/DLB | M | 78 | Unk | None |
| 25 | PSP | M | 72 | Unk | None |
| 26 | PSP | F | 86 | Unk | None |

We performed SV2C staining in substantia nigra and striatum (Fig. 8). We observed SV2C-positive staining in the SNpc and in the SNpr, as well as in terminal regions and cell bodies in the caudate nucleus and putamen. In PD, an unexpected and striking alteration in SV2C expression was revealed. We observed abnormal punctate SV2C-positive staining in the SNpc, and a similar pathology in the putamen (Fig. 8A) and caudate nucleus (not shown). This disruption in SV2C expression was found in each PD case, though it was less severe in the three cases with neuropathology consistent with comorbid DLB. Striatal SV2C staining was not altered in cases of AD, PSP, nor MSA with or without DLB comorbidity (Fig. 8B), suggesting a disruption in SV2C is a unique feature of PD.

**Figure 8.**
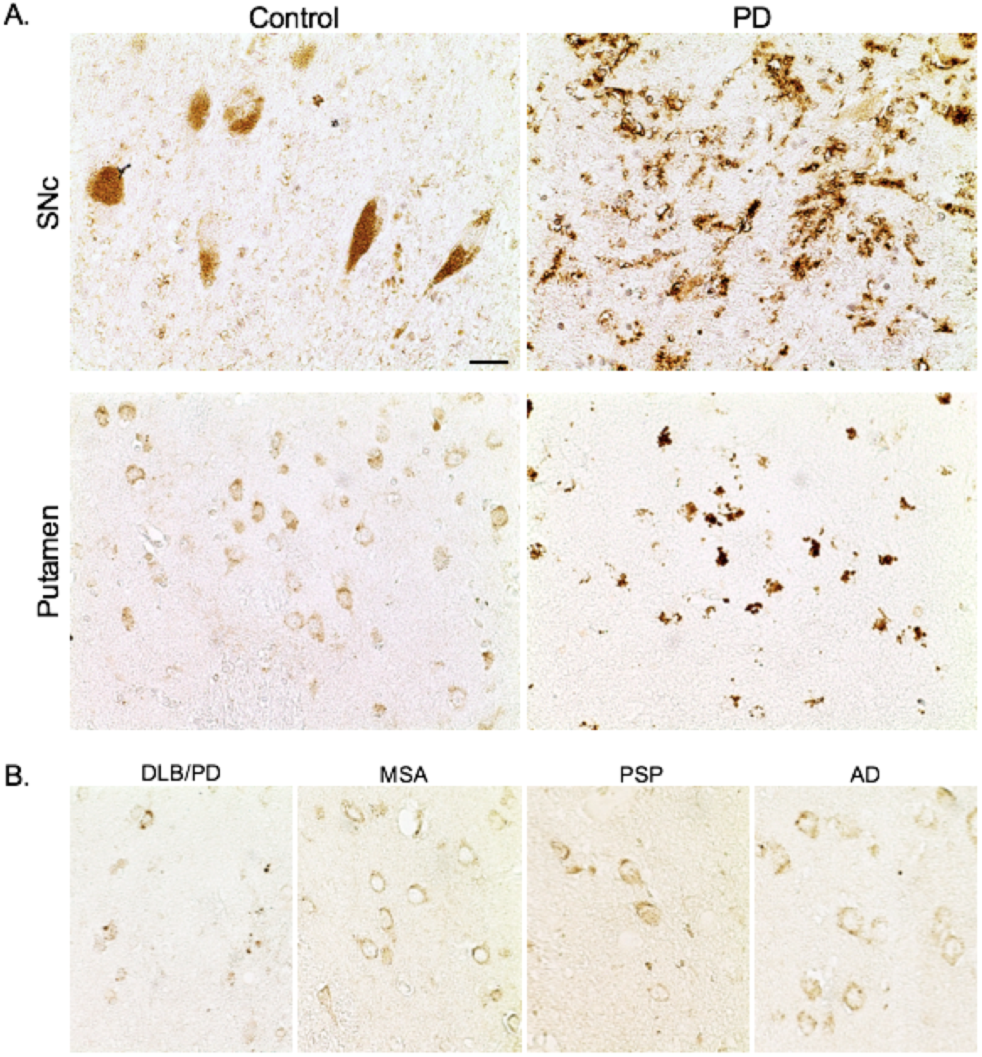
SV2C expression is specifically disrupted in PD basal ganglia. Representative micrographs demonstrate SV2C staining in human SNpc and putamen. (A) In PD, SV2C expression is significantly disrupted in the SNpc and putamen. SV2C-positive puncta are distributed throughout the SNpc and putamen in PD but not in age-matched controls.(E) Representative micrographs indicate that SV2C staining is relatively normal in the putamen of representative DLB/PD, MSA, PSP and AD cases. *Scale bar = 20μm*.

## DISCUSSION

The present data reveal that SV2C mediates dopamine dynamics and is disrupted in PD. We demonstrate that SV2C and α-synuclein interact, which may be important in disease pathogenesis. Further, we show that SV2C regulates dopamine release and dopamine content, as well as a neurochemical effect of nicotine, providing a molecular link to recent GWAS data identifying SV2C as a mediator of nicotine neuroprotection. Results from these experiments establish a novel role for SV2C and a basis for the potential functional consequence of a disruption of SV2C in PD and nicotine neuroprotection.

### Altered SV2C expression patterns in A53T α-synuclein overexpressors, but not other models of PD

As expected, striatal SV2C expression was slightly decreased following significant loss of dopaminergic terminals in mouse models of PD. Unexpectedly, SV2C-positive puncta were observed in the striatum of A53T α-synuclein OE animals. We did not observe similar punctate staining after MPTP lesion or in control animals: we modeled varying degrees of dopamine cell loss with multiple MPTP paradigms, including a moderate dose that closely matches the dopaminergic loss observed in A53T α-synuclein OE mice. This indicates that the observed disruption in SV2C is related to a distinct pathogenic mechanism that is not recapitulated by mere nigrostriatal dopaminergic degeneration. Unlike the A53T-OE model, MPTP administration in mice does not typically induce α-synuclein disruption(43), which further indicates that a link between α-synuclein and SV2C dysregulation may be particularly important. It is possible that SV2C pathology is not observed after MPTP intoxication due to the relatively acute nature of the lesion; however, we did not observe SV2C deposition in another genetic model lacking striatal α-synuclein aggregation, the VMAT2-LO mice, which suggests that even a life-long pathogenic mechanism leading to dopamine cell loss is not sufficient to spur similar SV2C disruption.

### SV2C-KO results in reduced dopamine tone and motor deficits

To further explore if any functional consequences of disrupted SV2C on the dopamine system in humans or animal models, we first needed to characterize involvement of the protein in basal ganglia functioning. SV2 proteins appear to promote vesicular function in various ways, perhaps in part by stabilizing transmitter content in the vesicle through neurotransmitter interaction with the protein’s heavily glycosylated intraluminal loop(32, 33) or by coordinating vesicular mobilization(24, 25, 44, 45) or by interacting with synaptotagmin-1 to facilitate calcium-stimulated exocytosis(26–30, 46, 47). Though only SV2A has been strongly implicated in epilepsy, recent evidence has hinted at a potential link between SV2C, dopamine and PD(37, 48, 49). Our neurochemical data directly indicate that SV2C plays a particularly important role in the dopaminergic nigrostriatal pathway. Here, we demonstrate that genetic deletion of SV2C results in a significant reduction in total dopamine content in the dorsal striatum; this dampened dopamine tone does not appear to result from aberrant dopamine metabolism, as the ratio of dopamine to its metabolites is unaltered. Pursuant to the observed reduction in total dopamine, genetic deletion of SV2C results in lowered release of dopamine evoked by electrical stimulation as measured by fast-scan cyclic voltammetry. Accordingly, we demonstrate that SV2C-KO animals have deficits in dopamine-related motor behavior, including reduced stride length and reduced locomotor activity. The reduction in locomotor activity observed in the SV2C-KO animals is primarily driven by apparent disruptions in circadian-mediated spikes of activity at the end of the wakeful period. This phenotype is consistent with other models of reduced dopamine vesicle function, such as the VMAT2-LO mice that exhibit motor impairment, reduced locomotor activity, altered circadian rhythm and smaller body size(39). Notably, SV2C-KO animals are not impaired on the rotorod test, a test which our lab previously demonstrated as insensitive in detecting motor deficits after moderate dopamine loss(50). Together, this behavioral and neurochemical data underscore that SV2C is indeed involved in basal ganglia performance and provides support for a potential functional and disease relevance for SV2C in PD.

### SV2C mediates a neurochemical effect of nicotine

Nicotine application normally results in a “high-pass filter” effect in which dopamine release is reduced upon low-intensity stimulations but enhanced upon high-intensity stimulations in the presence of 500nM nicotine. Nicotine, then, is thought to increase effects of salient dopaminergic inputs while dampening effects of transient dopamine release. Genetic deletion of SV2C ablates this effect of acute nicotine such that high-intensity stimulations elicit less dopamine than baseline release. This may indicate a functional interaction between SV2C and nicotine. The precise relationship between SV2C and nicotine neuroprotection remains unknown, but these data indicate that SV2C-KO animals may have altered neurochemical, behavioral, or neuroprotective response to nicotine. These data may hold relevance for recent GWAS data that identifies SV2C as a genetic mediator of the neuroprotective effects of nicotine.

### A link between altered SV2C expression and α-synuclein disruption

Another link between SV2C and PD highlighted by our data is an association between SV2C and α-synuclein. We demonstrate that the two proteins co-immunoprecipitate, indicating a physical and perhaps functional interaction. This interaction may be the basis for our observed mutual disruption of SV2C and α-synuclein in our mouse models. SV2C-KO animals have increased expression of high-molecular weight α-synuclein with commensurately decreased low-molecular weight α-synuclein. This is notable, as the ratio of monomeric to multimeric α-synuclein is likely important in the induction of α-synuclein aggregation and toxicity(51, 52). Our findings are supported by previous evidence indicating that SV2C and α-synuclein gene transcripts are normally highly correlated, but that this relationship is abolished in PD(53). These data suggest an association between SV2C and α-synuclein, and that disruptions in α-synuclein or SV2C may modify this interaction.

### SV2C disruption in Parkinson’s disease basal ganglia

Finally, we directly connect SV2C to human PD by demonstrating altered SV2C expression in PD basal ganglia. SV2C is normally distributed throughout the basal ganglia in dopaminergic regions of the SNpc and striatum in humans. This pattern of expression was largely unaltered in several neurodegenerative diseases affecting the basal ganglia, such as MSA, PSP, and DLB, as well as in non-basal ganglia-related AD. However, SV2C expression was consistently altered in PD: SV2C staining was almost entirely punctate in the nigrostriatal tract, revealing intra- and extracellular inclusions of SV2C. These data expose a novel pathologic feature not previously identified in PD. It is important to emphasize that this pathology appears to be largely restricted to PD. While basal ganglia pathologies are characteristic of PD, MSA, PSP and DLB, out of these, SV2C disruption is exclusive to PD, and is less severe in PD with neuropathology consistent with comorbid DLB. Though MSA, PSP and DLB are all classified as Parkinson’s Plus Syndromes, they differ clinically, etiologically and histopathologically from PD and are not effectively treated by dopamine replacement therapeutics (reviewed in Mitra, et al.(54)). The lack of SV2C pathology in PSP, DLB and MSA further distinguish them from PD and suggests that SV2C disruption is related to the concurrent α-synuclein disruption and dopaminergic degeneration characteristic of PD. It is unknown if similar SV2C disruption is present in forms of PD lacking Lewy body pathology, such as cases stemming from certain LRRK2 mutations(55–57). SV2C was not disrupted in AD, further emphasizing that SV2C disruption is not merely related to a neurodegenerative disease state. As with other neurodegenerative disease-related protein disruptions, more research is required to fully characterize the alteration in SV2C expression in PD; however, these data directly connect SV2C protein to PD for the first time and strengthen previous data implicating SV2C as a genetic mediator of PD risk in certain populations.

### A role for SV2C and PD pathogenesis

The cause or implications of SV2C deposition in the basal ganglia is unclear. As our mouse data suggest, SV2C is particularly important presynaptically in regulating dopamine release and in maintaining dopamine tone. Disruptions in SV2C may negatively affect dopaminergic vesicular function, as well as neuron integrity and neurotransmission, thereby contributing to human disease progression. As such, modulating the function of SV2C in order to enhance or restore its ability to preserve proper basal ganglia neurotransmission may be a promising avenue for therapeutics. Indeed, SV2C is a feasible target for pharmacotherapy: SV2C binds botulinum neurotoxin A(38, 58), and its close family member, SV2A, is the specific target for the antiepileptic drug levetiracetam(59). Though beyond the scope of this work, experiments in our laboratory are also underway to delineate the relative importance of SV2C in other neuronal populations. As our human data demonstrates additional localization of SV2C to striatal GABAergic cells, manipulating SV2C function may have functional and therapeutic implications for both dopamine and GABA neurotransmitter systems.

Finally, the severity and specificity of SV2C disruption in PD is striking. It suggests a novel pathologic feature unique to PD, which further distinguishes it from other basal ganglia-related neurodegenerative disorders. SV2C is localized to the basal ganglia, is associated with variable PD risk in smokers, promotes proper dopamine homeostasis and motor function, and is disrupted in PD. Taken together, these data establish a role for SV2C in nigrostriatal dopamine neurotransmission and identify it as a potential contributor to PD pathogenesis.

## MATERIALS & METHODS

### Antibodies

Two rabbit polyclonal anti-SV2C sera were raised against a peptide in the N-terminus region (amino acids 97–114) of SV2C: one against mouse SV2C (mSV2C; sequence STNQGKDSIVSVGQPKG), and one against human SV2C (hSV2C; sequence SMNQAKDSIVSVGQPKG). Peptides were conjugated to Imject^®^ Maleimide Activated mcKLH (Thermo Scientific) and sera were generated for our laboratory using Covance Custom Immunology Services. DAT and TH primary antibodies were purchased from Millipore, β-actin was purchased from Sigma. Polyclonal goat-anti SV2A (E-15) and SV2B (E-18) were purchased from Santa Cruz Biotechnology. Monoclonal mouse-anti SV2B and mouse-anti synaptotagmin-1 antibodies were purchased from Synaptic Systems. All secondary antibodies were purchased from Jackson ImmunoResearch Laboratories (biotinylated, HRP-conjugated) or Abcam (fluorescent)

### Tissue culture

HEK293 (ATCC) and Neuro-2a (N2a) cells were cultured according to standard protocols. Growth media was either DMEM with 10% fetal bovine serum (FBS) + 1% penicillin/streptomycin (HEK293) or EMEM + 10% FBS + 1% penicillin/streptomycin (N2a). Transfections of SV2C and SV2A in pcDNA3.1 vectors were performed with Lipofectamine 2000 (Invitrogen) according to manufacturer’s protocols. SV2C shRNA constructs were transfected via electroporation (Amaxa Nucleofector) according to manufacturer’s protocols. Cells were harvested 24 hours post-transfection, lysed in RIPA buffer and total protein extraction was achieved through differential centrifugation

### SV2C shRNA

Short hairpin RNAs (shRNA) against SV2C were custom-designed by Origene. Two SV2C-specific sequences (GACAGCATCGTGTCTGTAG and ATCGTGTCTGTAGGACAGC) were found to significantly reduce expression of endogenous SV2C expression in Neuro-2a cells

### Western blotting

Western blots were performed as previously described(60). Primary antibodies were used at the following dilutions: SV2C (1:2,500), DAT (1:5,000), TH (1:1,000), synaptotagmin 1 (1:1,000), α-synuclein (1:1,000), and β-actin (1:5,000). HRP-conjugated secondary antibodies were diluted to 1:5,000

### Animals

12-month old male and female α-syn OE mice and WT littermates, and 22-month old male and female VMAT2-LO mice and WT littermates were used for reported studies requiring genetic models of PD. Six- to 12-month old male WT C57BL/6 mice (Charles River Laboratories) were used for MPTP studies. Three to six animals were included in each group for all experiments. All procedures were conducted in accordance with the National Institutes of Health Guide for Care and Use of Laboratory Animals and previously approved by the Institutional Animal Care and Use Committee at Emory University

### MPTP treatments

Male mice were injected (s.c.) with either MPTP (Sigma) or saline. The “terminal” lesion consisted of two injections of 15 mg/kg MPTP (12-hr inter-injection interval). The 4x15mg/kg lesion consisted of 4 injections in 1 day of 15mg/kg MPTP (2-hr inter-injection interval). The “cell body” lesion consisted of five injections of 20 mg/kg MPTP (24-hr inter-injection interval)

### Transgenic mouse generation

#### SV2C-KO mice

We obtained 3 lines of C57BL/6 ES cells from the European Conditional Mouse Mutagenesis Program containing the SV2C “knockout-first construct”(61). ES cells containing the construct were injected into blastocysts, and we bred chimeric mice with C57BL/6 mice. We achieved germline transmission of the construct in the resulting offspring, and chose these mice to found the SV2C-“knockout first” line. We then bred this line with Flp-containing mice followed by mice containing Cre under a *NES*(nestin) promoter to delete the intron and found the SV2C-KO line. PCR genotyping primers are as follows: *Sv2c*(Exon 2): (Fw) TCA TCT AGA AGG GTT AAG GTC TGG, (Rev) ACC ATC ATC CCG AGG TAC AC; *LoxP*: (Fw) GCC TCA ACC AGA CCT AAG AA, (Rev) TAG GAA CTT CGG AAT AGG; *LacZ*: (Fw) GTC GTT TGC CGT CTG AAT TT, (Rev) CAT TAA AGC GAG TGG CAA CA; *Cre*: (Fw) CCT GGA AAA TGC TTC TGT CCG TTT GCC, (Rev) GAG TTG ATA GCT GGC TGG TGG CAG ATG; *Flp*: (Fw) CAC TGA TAT TGT AAG TAG TTT GA, (Rev) CTA GTG CGA AGT AGT GAT CAG G

#### Back-crossed VMAT2-LO mice

Mice expressing 5% of normal VMAT2 levels were generated as described previously(5, 10, 39) and back-crossed to a C57BL/6 genetic background(62). Mice were aged 22 months prior to immunohistochemical analysis

#### Alpha-synuclein overexpressing mice

Mice overexpressing human PD-associated α-synuclein A53T missense mutation in nigrostriatal dopamine cells under the control of a *Pitx3* promoter were generated as described previously(40). Animals were aged 12 months prior to immunohistochemical analysis

### Immunohistochemistry

#### Paraffin-embedded sections

Sections were deparaffinized in xylenes and rehydrated in decreasing concentrations of ethanol. Endogenous peroxidase was quenched with 3% H_2_O_2_, followed by antigen retrieval in citrate buffer (pH 6.0) at 95°C.Nonspecific antibody binding was blocked with 3% normal horse serum. Primary antibody was diluted to 1:2,500 (hSV2CpAb) or 1:1,000 (α-synuclein, DAT and TH). secondary antibody (1:200) signal was enhanced with an avidin-biotin complex (Vector Laboratories) and developed with a 3–3’ diaminobenzidine (DAB) reaction for approximately 45 seconds

#### Frozen sections

Mice were sacrificed by rapid decapitation or transcardial perfusion and the brains removed immediately placed in 4% paraformaldehyde for fixation. Brains were placed in 30% sucrose prior to sectioning. Brains were sectioned to 40µm. Staining was conducted as described previously(60, 63), except that all washes and dilutions were conducted in PBS with 0.2% Triton X-100

### Immunoprecipitation of SV2C and α-synuclein

Co-immunoprecipitation experiments were performed using the Pierce Co-Immunoprecipitation Kit (Thermo Scientific) according to manufacturer’s protocols. Briefly, mSV2CpAb was cross-linked to agarose beads. WT animals were sacrificed by rapid decapitation and a bilateral striatal dissection was performed followed by homogenization and centrifugation to achieve a synaptosomal preparation. Samples were treated with IP Lysis/Wash buffer (25 mM Tris-HCl pH 7.4, 150 mM NaCl, 1 mM EDTA, 1% NP-40 and 5% glycerol) incubated with antibody-bound resin overnight at 4°C and protein complexes were eluted with low-pH elution buffer

### Locomotor activity

Animals were placed in individual cages within locomotor activity monitoring boxes two hours prior to onset of dark (waking) period. Beam-breaks detected by accompanying software were recorded as a measure of total ambluations for a 24-hour observation period

### Gait Analysis

Animals were trained to walk along a clean paper path to their home cage. On testing day, forepaws were dipped in water-soluble ink and animals were prompted to transverse the paper. Stride length was measured as the total toe-to-toe distance across an average of 3 steps

### Rotorod

Animals were trained to balance on a slowly rotating rod. On testing day, animals were placed on the rod which accelerated to 12RPM over 10 seconds. Latency to fall, speed of rotation at the time of fall, and total distance traveled was recorded and averaged over three testing trials

### Fast-scan cyclic voltammetry in striatal slices

Fast-scan cyclic voltammetry was performed as described previously(60) in 300µm striatal slices bathed in 30°C oxygenated aCSF. Stimulations were 700µA 400ms monopolar pulse, either single pulse (baseline release) or five at 100Hz (high-intensity stimulation in acute nicotine experiments). Four to five recordings were taken at each of four dorsolateral striatal sites with a 5-minute inter-stimulation rest. Electrode sensitivity was calibrated to known dopamine standards using a flow-cell injection system. Maximum dopamine release was averaged across sites and kinetic constants were calculated using nonlinear regression analysis of dopamine release and uptake. *Acute nicotine application*: aCSF bath was replaced with increasing concentrations of dopamine: 10nM, 50nM, 100nM and 500nM for approximately 30 minutes each. Five recordings were taken at each concentration, with the exception of the 500nM concentration in which recordings were taken with five single-pulse stimulations and one 5 pulse/100Hz stimulation

### HPLC in striatal tissue

HPLC for striatal dopamine and DOPAC content was performed as described previously(64)

### Statistics

All data were analyzed in GraphPad Prism. Differences between genotypes or treatments were compared by two-tailed t tests. Densitometric analyses were performed with Image Lab software. All errors shown are SEM, and significance was set to *p < 0.05*

## Acknowledgements

The authors would like to thank Dr. Marla Gearing for her expert insight and Merry Chen for excellent technical assistance. This research project was supported by NIEHS R01ES023839 and NIEHS P30ES019776 to G.W.M., NINDS F31NS089242 to A.R.D.,the Neuropathology Core of the Emory Neuroscience NINDS Core Facilities (NIH P30NS055077), The University of Washington ADRC Brain Bank (NIH P50AB005136), the Lewis Dickey Memorial Fund, and the intramural research programs of the NIH (NIA AG000928 and AG000929) to H.C

## Authorship

Conceptualization, A.R.D., K.A.S., K.M.L., W.M.C., and G.W.M. Methodology, A.R.D., K.A.S.,K.M.L., A.I.B, Y.L., M.W., H.C., C.S., N.S. Investigation, A.R.D., K.A.S., M.O., K.M.L. Resources, H.C. Formal analysis, A.R.D. and K.A.S. Interpretation of results, A.R.D., M.O., W.M.C., K.A.S., H.C., and G.W.M. Writing—original draft, A.R.D. and G.W.M. Writing— review and editing, all authors. Supervision, W.M.C, H.C.,G.W.M. Funding acquisition, A.R.D., H.C., G.W.M

